# Spontaneous Blink Rate Correlates With Financial Risk Taking

**DOI:** 10.1101/046821

**Authors:** Emily Sherman, Robert C Wilson

## Abstract

Dopamine has long been thought to play a role in risky decision-making, with higher tonic levels of dopamine associated with more risk seeking behavior. In this work, we aimed to shed more light on this relationship using spontaneous blink rate as an indirect measure of dopamine. In particular we used video recording to measure blink rate and a decision-making survey to measure risk taking in 45 participants. Consistent with previous work linking dopamine to risky decisions, we found a strong positive correlation between blink rate and the number of risky choices a participant made. This correlation was not dependent on age or gender and was identical for both gain and loss framing. This work suggests that dopamine plays a crucial and quite general role in determining risk attitude across the population and validates this simple method of probing dopamine for decision-making research.

## Introduction

Many decisions in daily life are made under considerable uncertainty. This uncertainty comes in many forms, from the well-defined risks of playing roulette, to the more ambiguous odds that it will rain on your wedding day. Across the population, there is a large amount of variability in how we deal with this uncertainty (e.g. Weber et al., 2002; Tymula et al. 2013) and there is considerable interest in understanding how these individual differences arise.

One factor that is thought to play a crucial role in risk taking is the neurotransmitter dopamine. In particular dopamine is thought to be important for determining our attitudes toward risk, the kind of uncertainty that arises when the odds of winning are known (e.g. in a game of roulette). For example, dopamine-related genes have been associated with risk taking (Kuhnen & Chiao, 2009; Dreber et al. 2009; Dmitrieva et al. 2010; Farrell et al. 2012) and dopamine drugs have been found to modulate risky behavior in both humans and animals (Riba et al. 2008; St Onge & Floresco, 2009; Norbury et al. 2013; Rutledge et al. 2015; although see Symmonds et al. 2013). Moreover, prolonged treatment with L-DOPA (a dopamine agonist) in Parkinson’s is associated with pathological gambling in a subset of patients (Molina et al. 2000; Santangelo et al. 2013).

Despite this progress, the exact nature of the relationship between dopamine and risk taking is incompletely understood. For one thing, dopamine has different effects on different receptors, which are themselves distributed differently in different areas of the brain (reviewed in Hurley & Jenner, 2006). Moreover, some studies have found that dopamine genes and drugs have different effects depending on range of other factors including gender (Dreber et al. 2009), baseline sensation seeking (Norbury et al., 2013) and whether the gambles involve gains, losses or a mixture of the two (Rutledge et al., 2015).

In this work we sought to shed more light on dopamine’s role in risk taking by using a remarkable relationship between the rate at which someone blinks and the amount of dopamine in their brains (Karson, 1983; Karson, 1988; Lawrence & Redmond 1991; Kleven & Koek, 1996). In particular, more frequent blinking is associated with greater dopamine in the striatum, a relationship that appears to be dependent on D2 (and possibly D1) receptors (Elsworth et al., 1991; Jutkiewicz and Bergman, 2004). We therefore hypothesized that if blink rate reflects dopamine and dopamine drives risk taking, then we should see a positive relationship between individual differences in spontaneous blink rate and risk taking across the population. By including age, gender and gambles of different valence, we also aimed to test whether the relationship between dopamine and risk was modulated by these factors as predicted by previous work.

## Methods

### Participants

Forty five adults (17 male, 28 female of which 21 were students all aged 18, and 24 were older, ages 26-59) were recruited from the students, teachers and parents at BASIS school in Phoenix. All subjects gave informed consent and the study was approved by the institutional review board at BASIS.

### Experiment

Each participant was seated in a quiet room judged to be quiet and lacking distractions. The participant was given the consent form to read and then sign. Then the details of what was expected of the participant were carefully explained by the experimenter (ES). The experiment itself consisted of two parts, first measurement of spontaneous blink rate and second a risk-taking survey.

#### Measurement of blink rate

We measured spontaneous blink rate by recording a movie of the participant while they “stared into space”. The movie was recorded on the webcam of an Apple laptop computer that was placed on the table in front of the participant. Participants were told to look straight ahead for two minutes while we filmed them and they were instructed to act as normally as possible during this period. They were informed that we were filming them but were not told that we were measuring their blinks. Blink rates were then computed manually by the experimenter while the participants completed the decision making survey. To ensure privacy for the participant, the video was deleted in front of the participant at the end of the experiment. All other data is available in the Supplementary Material (S1 Data) along with code we used to process it (S1 Code).

#### Decision-making survey

Once the two minutes had passed, the webcam of the computer was shut off and the participant was handed a paper survey. The survey consisted of nine questions, with each question offering participants a choice between a certain outcome (e.g. 100% chance of $240) and a risky outcome (e.g. 25% chance of $1000). For each question, participants had to choose which option they would prefer. The gambles were only ever hypothetical and participants were not paid for their time or on the basis of their choices.

The questions themselves were chosen based on the results of a pilot study conducted at the University of Arizona that had revealed a possible relationship between blink rate and answers to these questions. In particular the nine questions were:

1. If you were faced with the following choice which alternative would you choose?
  a. A sure gain of $240
  b. A 25 percent chance to gain $1000 and a 75 percent chance to gain nothing.
2. If you were faced with the following choice which alternative would you choose?
  a. A sure loss of $750
  b. A 75 percent chance to lose $1000 and a 25 percent chance to lose nothing?
3. In addition to whatever you own you have been given $2000. You are now asked to choose between:
  a. A 50% chance of losing $1000
  b. A sure loss of $500
4. If you were faced with the following choice which alternative would you choose?
  a. A 100 percent chance of losing $50
  b. A 25 percent chance of losing $200 and a 75 percent chance of losing nothing
5. If you were faced with the following choice which alternative would you choose?
  a. A sure loss of $3000
  b. An 80 percent chance to lose $4000 and a 20 percent chance to lose nothing.
6. If you were given a choice which of the following gambles would you prefer?
  a. $1,000,000 for sure
  b. A 10 percent chance of getting $2,500,000, an 89 percent chance of getting $1,000,000 and a 1% chance of getting $0
7. In addition to whatever you own you have been given $1000. You are now asked to choose between:
  a. A 50% chance of getting $1000
  b. A sure gain of $500
8. If you were faced with the following choice which alternative would you choose?
  a. A sure gain of $3000
  b. An 80 percent chance to gain $4000 and a 20 percent chance to gain nothing
9. Suppose you are offered the chance to play the following game: I flip a fair coin. If it comes up heads you lose $100. If it comes up tails you win $125. Do you accept?
  a. Yes
  b. No

## Results

### Distribution of blink rates is consistent with earlier findings

Across the population we observed a mean blink rate of 21 blinks per minute, a finding which is consistent with the literature (e.g. Bentivoglio et al., 1997). In line with earlier findings we also found a wide distribution of blink rates across the population (Figure 1A). Breaking out results for gender and age (treated as a discrete variable for younger, age < 19, and older), there was a numerical hint of an interaction such that young male participants blinked more frequently than other groups, although this was not statistically significant (2x2 ANOVA with age group (young/old) and gender as factors: age F(1,44) = 1.03, p = 0.32; gender F(1,44) = 1.65, p = 0.21; age × gender F(1,44) = 0.8, p = 0.38) (Figure 1B). Treating age as a continuous variable in a linear regression model with age, gender and the age × gender as factors, gave similar results (β(age) = −0.10, p = 0.36; β(gender) = −4.03, p = 0.29; β(age×gender) = 0.06, p = 0.58).

**Figure 1.**
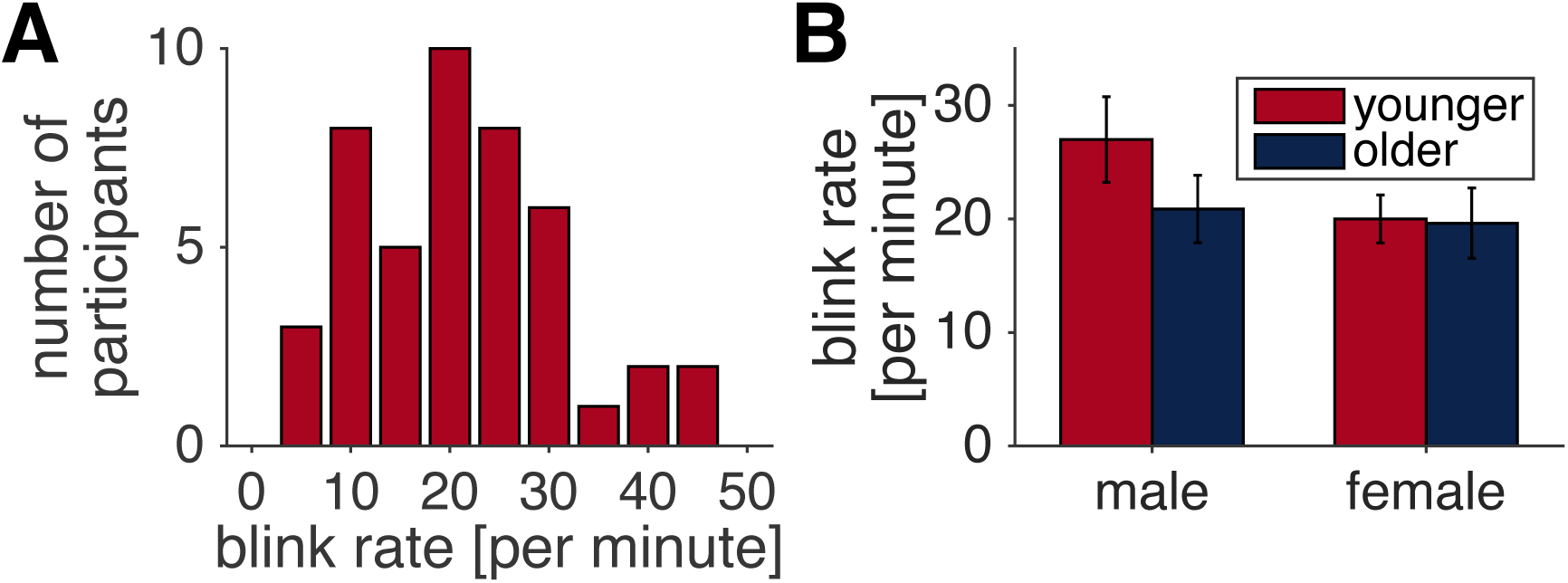
Blink rates across the population. (**A**) Distribution of blink rates across the population. (**B**) Blink rates are numerically higher for young male participants than any other group, although this difference is not statistically significant.

### Distribution of risk seeking across the population

In our simplest measure of risk seeking, we counted the number of risky choices (out of 9) made by each participant. As with blink rate, there was a large range across the population (Figure 2A). Unlike blink rate we found a weak main effect of age on blink rate when treating age as a discrete variable, such that older participants were found to blink less than younger adults (2x2 ANOVA with age and gender as factors, age F(1,44) = 4.17, p = 0.05; gender F(1,44) = 1.91, p = 0.17; age x gender F(1,44) = 1.58, p = 0.22). This effect of age seemed to be stronger for men than for women (although the interaction was not significant) and post hoc t-tests suggest a trend level effect for men but not women (for men, two-sided t-test, t(19) = 2.00, p = 0.06; for women t(26) = 0.65, p = 0.52). Treating age as a continuous variable in a regression yielded similar results, although the significance of the age effect was reduced (β(age) = −0.03, p = 0.10; β(gender) = −0.75, p = 0.22; β(age×gender) = 0.01, p = 0.50)

**Figure 2.**
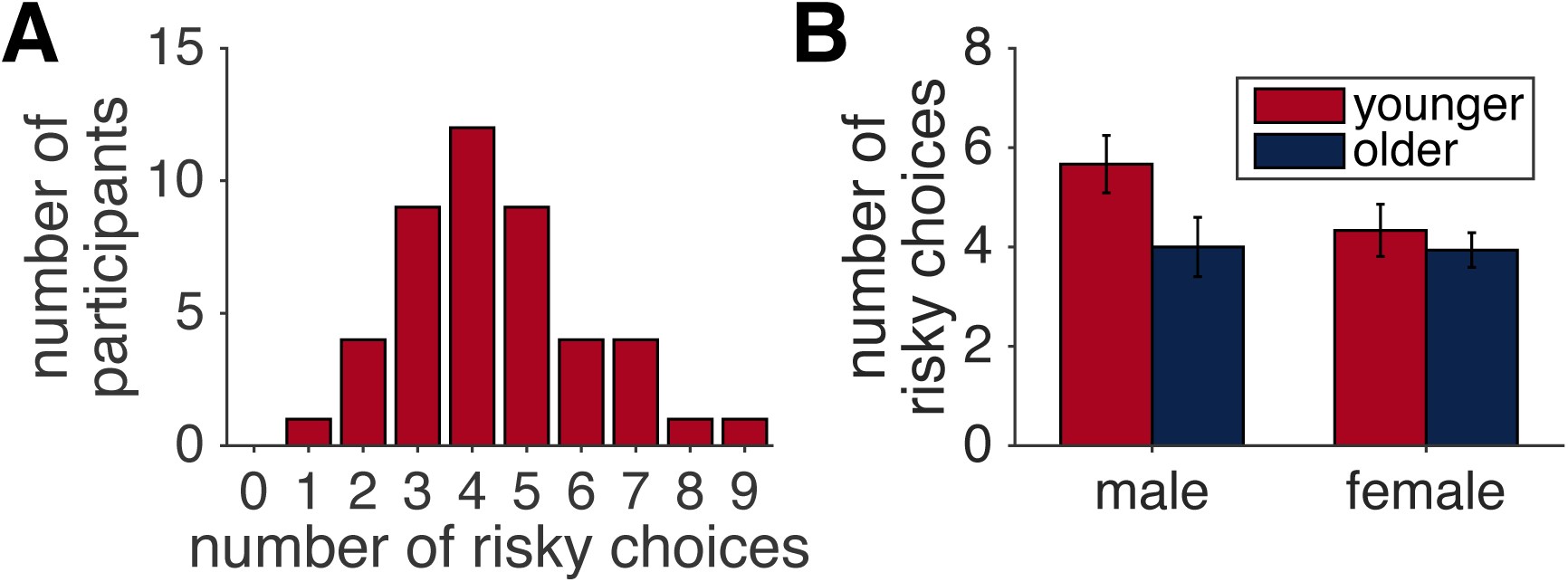
Risk preference across the population. (**A**) Distribution of risk seeking across the population. (**B**) Risk seeking declined with age for male, but not female, participants.

We also looked at behavior on the individual items. This revealed a relatively wide range of preferences across the questions, from about 30% of people choosing the risky option in question 1 to about 60% choosing the risky option in question 3 (Figure 3A). Given the variation in expected value of the gambles in our questions, this wide range of behavior was not unexpected. In line with classic findings from the literature (Kahneman & Tversky, 1979), we also found risk seeking to be greater for the loss questions than the gain questions (Figure 3B, t(44) = −2.75, p = 0.009). However, it is important to note that (with the exception of questions 3 and 7) the questions did not equate expected value between gain and loss domains so it is important not to over interpret this result.

**Figure 3.**
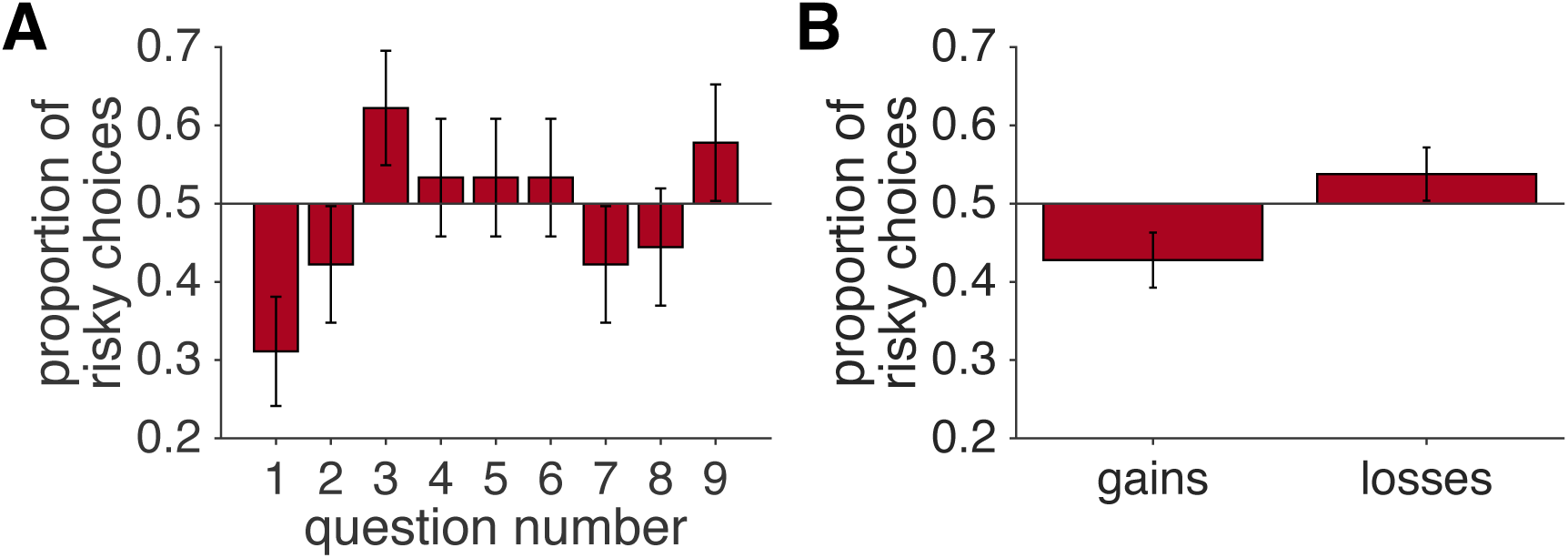
Item analysis. (**A**) The proportion of risky choices varied by about 30% across the nine different questions. (**B**) In line with classic findings, participants were more risk averse for gains than losses, although differences in expected value between gains and loss questions make it difficult to draw strong conclusions.

### Blink rate is positively correlated with risk seeking

In the most straightforward test of our hypothesis, we computed the correlation between the blink rate and the number of questions in which participants selected the risky option. This revealed a strong positive correlation between blink rate and risk seeking, such that participants with higher blink rates chose the risky option more frequently (Spearman’s ρ(43) = 0.57, p = 4.45 × 10^−5^) (Figure 4). This correlation also survives correction for age (treated continuously), gender and the interaction between age and gender, which we achieved by regressing out the effects of age, gender and the interaction on both blink rate and risk seeking (Spearman’s ρ(43) = 0.52, p = 2.6 × 10^−4^).

**Figure 4.**
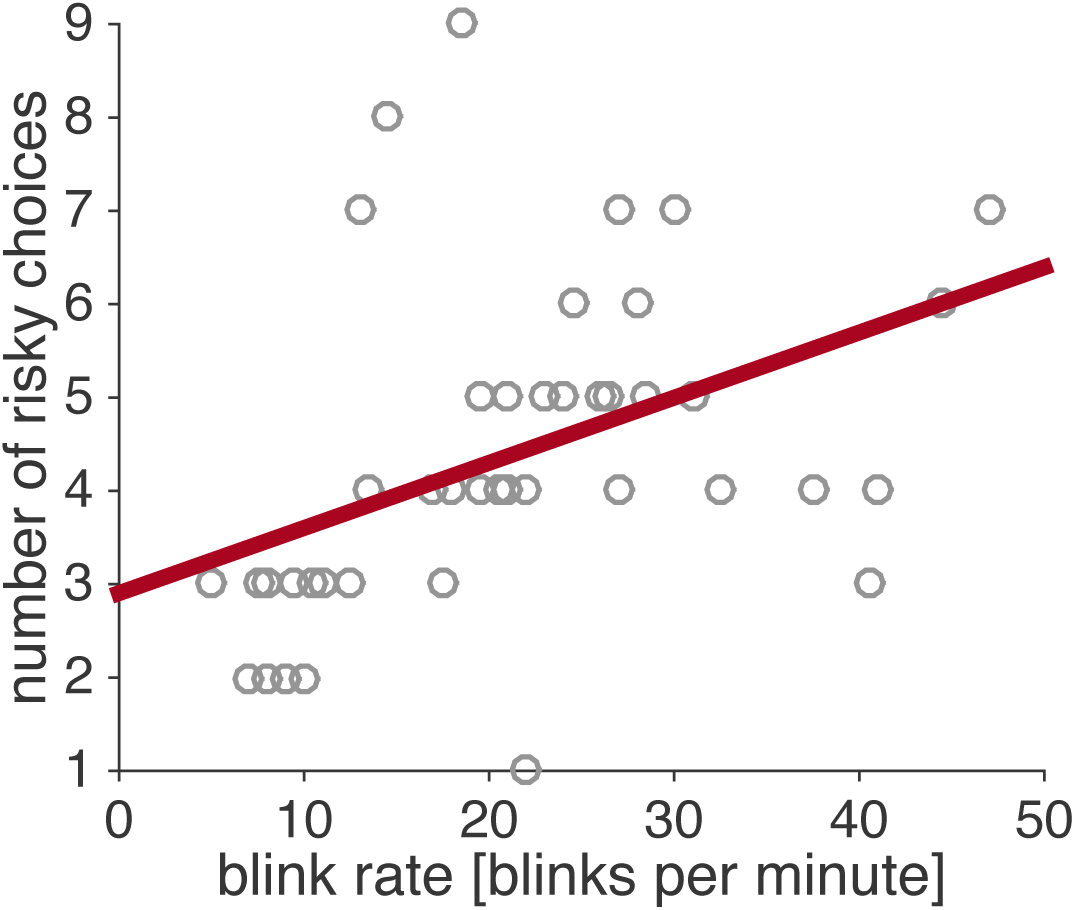
Individual differences in blink rate correlate with individual differences in risk seeking.

### Item analysis reveals effect is independent of gains and loss framing

To quantify the effect of blink rate on the choices of individual questions we turned to logistic regression. In particular we modeled the probability of choosing the risky option on each question as

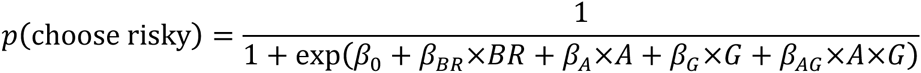

where *BR* is the blink rate, *A* is the age group (−1 for young, +1 for old), *G* is gender (-1 for male, +1 for female). The regression coefficients (*β*_0_, *β*_*BR*_, *β*_*A*_, *β*_*G*_ and *β*_*AG*_) were fit separately for each question. In figure 5A we plot the blink rate coefficient, *β*_*BR*_, for each of the nine different questions. While this regression weight approaches significance only for the last question (p = 0.06), it is interesting to note that, numerically, the size of these coefficients is similar for all questions and the sign is positive for all but one, suggesting that the same relationship between blink rate and risky decision making holds for all questions. In addition, there is no obvious difference between the coefficients for gain and loss questions, suggesting that blink rate modulates risk seeking regardless of valence. This is further illustrated in Figure 5B, in which we plot the proportion of risky choices for gain and loss questions against blink rate. This reveals a positive correlation for both gains and loss framing, the slope of which is nearly identical in the two cases (for gains, Spearman’s ρ(48) = 0.43, p = 0.004; for losses ρ(48) = 0.40, p = 0.007).

**Figure 5.**
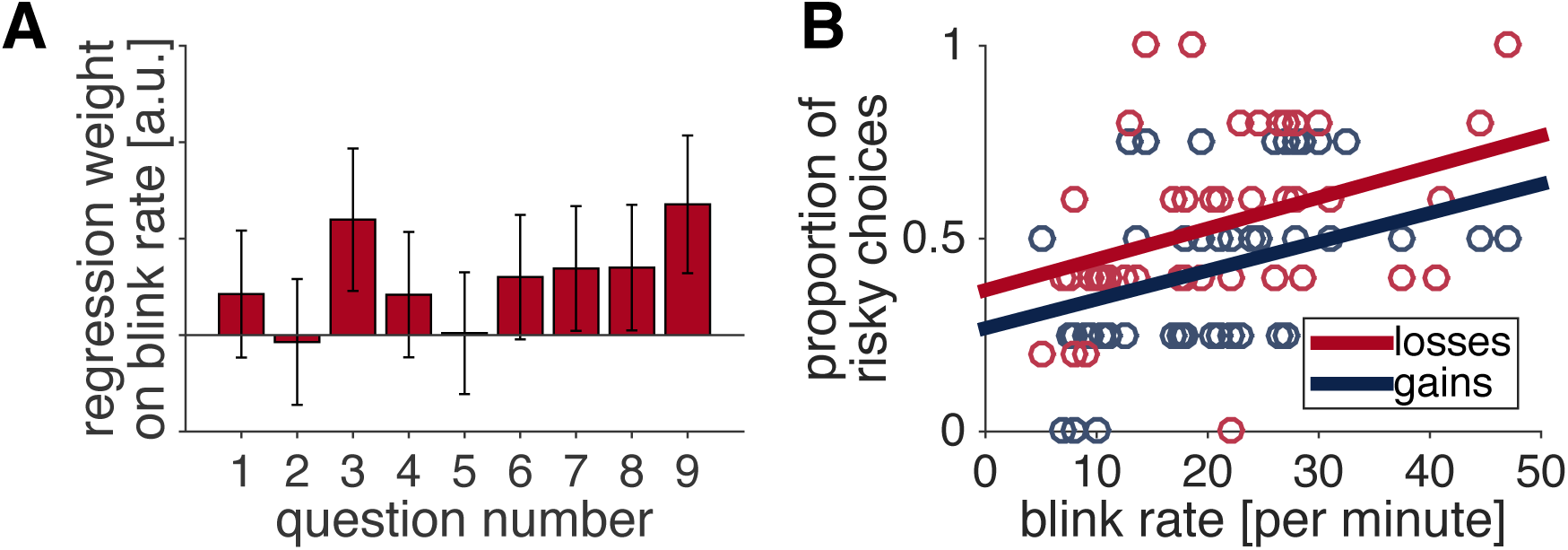
Similar relationship between blink rate and risk seeking for all questions. (**A**) Logistic regression yields a similar weight on blink rate for all nine questions. (**B**) Nearly identical relationship between blink rate and risk seeking in both gains and losses domains.

## Discussion

In this work we investigated the relationship between blink rate, a known measure of dopamine (e.g. Karson 1983), and risky decision making in a sample of 45 participants ranging in age from 18 to 59. Our findings suggest a strong relationship between blink rate and risk taking that is independent of age and gender and does not appear to be modulated by the valence of the decision, i.e. whether the risky choice is for gains or losses. This suggests a relatively general role for dopamine in promoting risk seeking over and above any biases induced by age, gender or framing of the problem.

The fact that we found almost identical relationships between blink rate and risk seeking for both gains and losses contrasts with recent work by Rutledge et al. (2015). In particular, these authors found that L-DOPA increases risk seeking only for gambles involving gains, but not for gambles involving losses or a mixture of losses and gains.

While there are many differences between the two experiments, two possibilities would be particular important to test. First is the difference in payoff structure. In our task the rewards were hypothetical and participants were not paid, while in the Rutledge task, participants were paid based on their choices. Thus participants in our experiment may not have taken the choices as seriously as the participants playing for real money and may have behaved differently as a result. Second is the different types of dopamine under consideration in the two experiments. In particular, blink rate has been associated with D2 (and possibly D1) related dopamine in striatum (Elsworth et al., 1991; Jutkiewicz and Bergman, 2004), while L-DOPA increases dopamine in a non-specific manner. It may therefore be the case that L-DOPA-related increases dopamine at other receptors and in other brain areas may counteract the effects of increased D2 activity in the Rutledge experiment. Combining blink rate and drug manipulation in a single study would be a first step to resolving this differences.

One limitation of our study is the relatively small number of questions we asked our participants. This was partly by design so that the experiment would be simple to run, however this limited number of questions makes it impossible to assess whether there is any interaction between blink rate, risk taking and the *quantitative* properties of the gambles themselves. Such interactions include the effects of reward magnitude, probability and outcome variance for the risky gamble are known to impact risk seeking (Kahneman & Tversky, 1979). Indeed, recent work by Norbury and colleagues has suggested that D2-and/or D3-related dopamine may play a role in how such quantitative properties of the gambles affect choice (Norbury et al. 2013).

In addition to the association with blink rate, we also found age-related differences in the risk-taking behavior of men, with young men taking more risk than older men. While this trend was similar to the numerical changes we saw in the blink rates of younger and older men, this numerical effect of age on blink rate was not significant, suggesting that changes in the risk attitude of men with age is not mediated by blink rate. This was slightly surprising given the well-known drop off in both dopamine level and receptor availability with age (Volkow et al., 1996). However, it is possible that other age-related changes in blinking, related to dryness of the eyes and mechanical changes to the eyelid, could mask any dopamine related changes in blink rate with age (Sun et al., 1997). Clearly more work will be needed to probe whether changes in dopamine drive changes in risk attitudes with age.

An obvious question for future work is whether our findings for decision-making under risk apply to other kinds of decision-making under uncertainty? For example, does blink rate correlate with decisions under ambiguity, in which the odds of the gamble are not known, in the same way that it correlates with risk? Previous work has suggested that risk and ambiguity preference are not correlated with one another across the population (Tymula et al., 2013) and may be driven by different neural processes (Hsu et al., 2005; Huettel et al. 2006). It may therefore be the case that ambiguity seeking does not correlate with blink rate in the same way that risk seeking appears to. Another example of particular relevance is that of decisions involving other kinds of risk, such as drug taking and sexual risk taking. Previous work has shown that risk preference can be highly domain specific (Weber et al., 2002) and it would be interesting to see whether blink rate correlates with risky behavior across decision-making domains.

## Acknowledgements

We would like to thank XXX at BASIS Scottsdale for helpful comments on the design and execution of this study.

## Supporting information captions

**S1 Dataset – Full dataset.** Including choices on individual questions, blink rate, gender and age.

**S1 Code – Matlab code reproducing all analyses and figures.**

